# A computational protocol to characterize elusive Candidate Phyla Radiation bacteria in oral environments using metagenomic data

**DOI:** 10.1101/358812

**Authors:** Peiqi Meng, Chang Lu, Xinzhe Lou, Qian Zhang, Peizeng Jia, Zhimin Yan, Jiuxiang Lin, Feng Chen

## Abstract

Several studies have documented the diversity and potential pathogenic associations of organisms in the human oral cavity. Although much progress has been made in understanding the complex bacterial community inhabiting the human oral cavity, our understanding of some microorganisms is less resolved due to a variety of reasons. One such little-understood group is the candidate phyla radiation (CPR), which is a recently identified, but highly abundant group of ultrasmall bacteria with reduced genomes and unusual ribosomes. Here, we present a computational protocol for the detection of CPR organisms from metagenomic data. Our approach relies on a self-constructed dataset comprising published CPR genomic sequences as a filter to identify CPR sequences from metagenomic sequencing data. After assembly and functional prediction, the taxonomic affiliation of CPR contigs can be identified through phylogenetic analysis with publically available 16S rRNA gene and ribosomal proteins, in addition to sequence similarity analyses (e.g., average nucleotide identity calculations and contig mapping). Using this protocol, we reconstructed two draft genomes of organisms within the TM7 superphylum, that had genome sizes of 0.594 Mb and 0.678 Mb. Among the predicted functional genes of the constructed genomes, a high percentage were related to signal transduction, cell motility, and cell envelope biogenesis, which could contribute to cellular morphological changes in response to environmental cues.

**Importance:** Candidate phyla radiation (CPR) bacterial group is a recently identified, but highly diverse and abundant group of ultrasmall bacteria exhibiting reduced genomes and limited metabolic capacities. A number of studies have reported their potential pathogenic associations in multiple mucosal diseases including periodontitis, halitosis, and inflammatory bowel disease. However, CPR organisms are difficult to cultivate and are difficult to detect with PCR-based methods due to divergent genetic sequences. Thus, our understanding of CPR has lagged behind that of other bacterial component. Here, we used metagenomic approaches to overcome these previous barriers to CPR identification, and established a computational protocol for detection of CPR organisms from metagenomic samples. The protocol describe herein holds great promise for better understanding the potential biological functioning of CPR. Moreover, the pipeline could be applied to other organisms that are difficult to cultivate.

## Introduction

The human oral cavity is one of the five primary microbial microecosystems within humans, and has been used as a model system for microbiome research (1). Dysbiosis of the oral microbial community has been observed in relation to some common systemic diseases including rheumatoid arthritis (2) and type 2 diabetes mellitus (3). Until recently the study of the oral microbiota has primarily focused on bacteria due to their relative high abundance, easy detection, and cultivability. In contrast, the ecological contributions of oral fungi, viruses, phages, and the newly classified “candidate phyla radiation” (CPR) are not fully studied. However, some of these microorganisms are also associated with diseases. For instance, *Candida albicans* is the most common fungi in the human oral cavity and has been implicated as the main etiological factor of oral candidiasis, which is universally developed in malignant tumors and HIV-infected patients (4). Further, bacteria within the TM7 division, which belong to the CPR group, have been reported to play an important role in multiple mucosal diseases including periodontitis, halitosis, and inflammatory bowel disease (5–8).

CPR lineages have been predicted to comprise greater than 15% of the domain Bacteria, and their previous identification represents an expansion of the tree of life (9). CPR organisms appear to share a number of characteristics. First, they have very limited biosynthetic and metabolic capabilities, with the absence of electron transport chains, tricarboxylic acid cycles, amino acid and membrane biosynthesis pathways, and various ribosomal subunits (9–14). Second, they have reduced genomes (often <1 Mb) (9), with the apparent absence of genes encoding CRISPR/Cas bacteriophage defense-systems (15). High-resolution cryo-TEM of CPR cells have revealed extremely small cell sizes of 0.009 ± 0.002 mm^3^, which is consistent with their small genome sizes (12). Taken together, these shared properties suggest that members of the CPR may exhibit symbiotic lifestyles, and are dependent upon partner cells for necessary metabolites, while potentially providing waste products from labile fermentation (e.g., acetate) in return (9, 16–18). Such co-dependent lifestyles could potentially account for their recalcitrance to *in vitro* cultivation (19).

Challenges in cultivating CPR organisms have limited the study of their phylogenetic and functional diversity. Further, 16S ribosome RNA (16S rRNA) gene-based detection methods are also not adequate for the classification of CPR species, due to the divergence of CPR 16S rRNA sequences and the inadequacy of PCR primers commonly used for gene amplification (9, 20). Moreover, their unusual 16S rRNA gene sequences, particularly those with unexpected insertions, may be discarded as artefacts (9). Lastly, many CPR lineages that are only known from 16S rRNA-targeted sequencing lack genomic representatives, which hinders the prediction of their potential functional significance.

With the development of high-throughput sequencing and bioinformatics analyses, increasing metagenomic sequencing data have been acquired from diverse ecological niches and multiple human body sites, including the human oral cavity. Metagenomics is a shotgun sequencing-based DNA sequencing methodology in which DNA isolated directly from the environment is sequenced, and the reconstructed genome fragments are assigned to draft genomes (21). This polymerase chain reaction (PCR)-independent genome- (rather than gene) based approach is of great value for researchers in overcoming the aforementioned characterization barriers, and provides information about the metabolic potential of uncultivated organisms, in addition to a variety of phylogenetically informative sequences that can be used to classify CPR organisms.

Here, we provide a computational protocol for discovering and characterizing CPR species. This protocol leverages metagenome shotgun sequencing data that has been acquired from the saliva of healthy controls and patients with the common mucosal disease, oral candidiasis.

## Results and Discussion

### Overview of metagenomic sequencing of salivary samples

Metagenomic shotgun sequencing was conducted on 26 salivary samples including 14 from oral candidiasis patients, and 12 from healthy persons. A total of 154.4 gigabases (Gb) of paired-end sequence data was generated, with an average of 27.6 million reads (5.5 Gb) per sample. After filtering low-quality reads, 97.2 ± 0.9% of the sequence reads remained. Human DNA accounted for 48 ± 23.2% of the high-quality reads and was also filtered prior to further processing (**Table S1**).

In the remaining, quality-filtered libraries, the percent of the reads mapped to the CPR database ranged from 0.12% to 1.51%, with no significant difference in abundance between the disease and control groups. There were also no significant differences in the abundance of reads mapped to single reference genomes for most of the CPR genomes in the database between the groups. The exceptions were *Candidatus* TM7_SBR2 and SBR3, which were found in a wastewater treatment plant through metagenomic sequencing, and the Candidate divisions RAAC1_SR1_1 and RAAC3_TM7_1, both of which were identified from deep metagenomic sequencing of microbial communities in acetate-stimulated aquifer sediments (**Fig. S1**).

Reads that mapped to the CPR reference database were merged separately in the disease and control groups, and then assembled into contigs. In the control group, the draft genome comprised 343 contigs with a total length of 0.594 Mb, and an average GC content of 48.0%. A total of 650 protein-coding sequences and a 16S rRNA gene with a length of 1178 bp were predicted in this genome. Likewise, 357 contigs were assembled into a draft genome within the disease metagenome group, comprising a total size of 0.678 Mb, and a GC content of 50.9%. A total of 1131 predicted protein-coding sequences were detected along with a single 16S rRNA gene with a length of 1273 bp in this genome.

### The CPR organisms in both the control and disease groups are members of the TM7 superphylum

Phylogenetic analyses of the 16S rRNA genes of the draft genomes constructed here along with previously reported sequences from CPR genomes (9, 33) and sequences in the SILVA database (30), confirmed that both of the CPR draft genomes identified in the disease and control groups belonged to the candidate division TM7 superphylum (**Fig. 1**). The 16S rRNA gene sequences in the control and disease groups shared 95% and 94% identity with their closest sequences in NCBI, respectively, both of which are from uncultivated clones (*Candidatus* TM7_s1 and *Candidatus* TM7_s7, respectively) from human oral gingival crevices.

**Fig 1.**
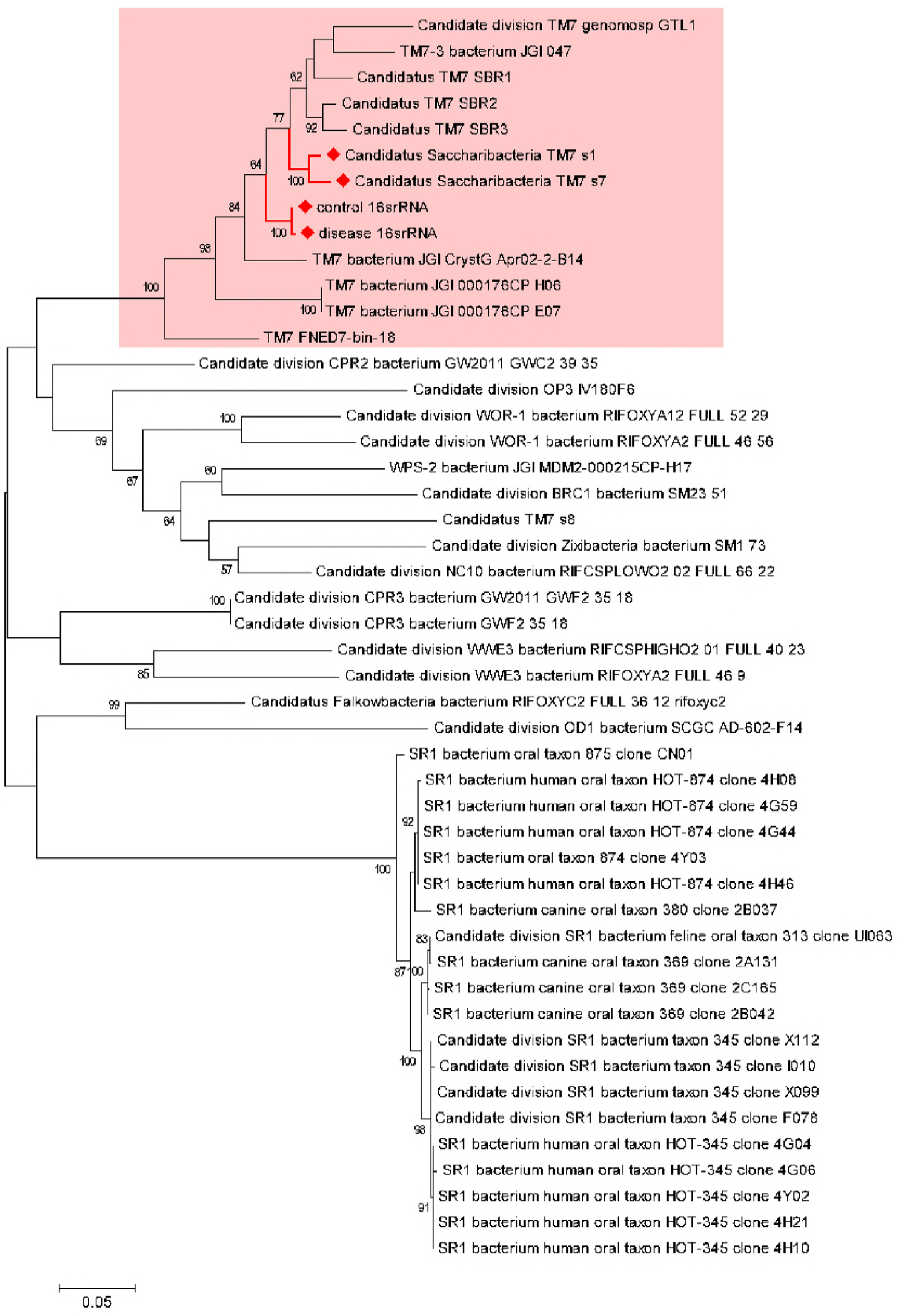
Maximum likelihood phylogeny of 16S rRNA genes from two newly constructed CPR draft genomes and previously published CPR genome representatives. Bootstrapping was conducted with 100 replicates and bootstrap values larger than 50 are indicated at the nodes. The newly identified CPR organisms and their closest relatives are indicated with red diamonds. The radiation of the TM7 superphylum is indicated by a pink background.

To further assess the phylogenetic positions of the newly identified CPR organisms, we aligned, concatenated, and phylogenetically analyzed 16 ribosomal protein sequences from each organism. This approach can yield higher-resolution trees and better resolve deeper divergences than that which is possible with single genes, including the widely used 16S rRNA gene (34). Further, the use of ribosomal proteins avoids artefacts arising from phylogenies constructed with genes with unrelated functions that are subject to different evolutionary processes. Moreover, ribosomal proteins are generally syntenic and co-located in a small genomic region in bacteria, reducing errors in phylogenetic analyses arising from spurious binning that could substantially perturb tree topologies. Finally, another important advantage of ribosomal protein trees compared to 16S rRNA gene trees is that they can include organisms with incomplete or missing 16S rRNA gene sequences (33). In the ribosomal protein-based phylogenetic tree, the CPR organisms of the control and disease groups belonged to the TM7 phylum lineage and were most closely related to *Candidatus* TM7_s1 and *Candidatus* TM7_s7 (**Fig. 2** and **Fig. S2**), which is in accordance with the 16S rRNA-based phylogenetic tree.

**Fig 2.**
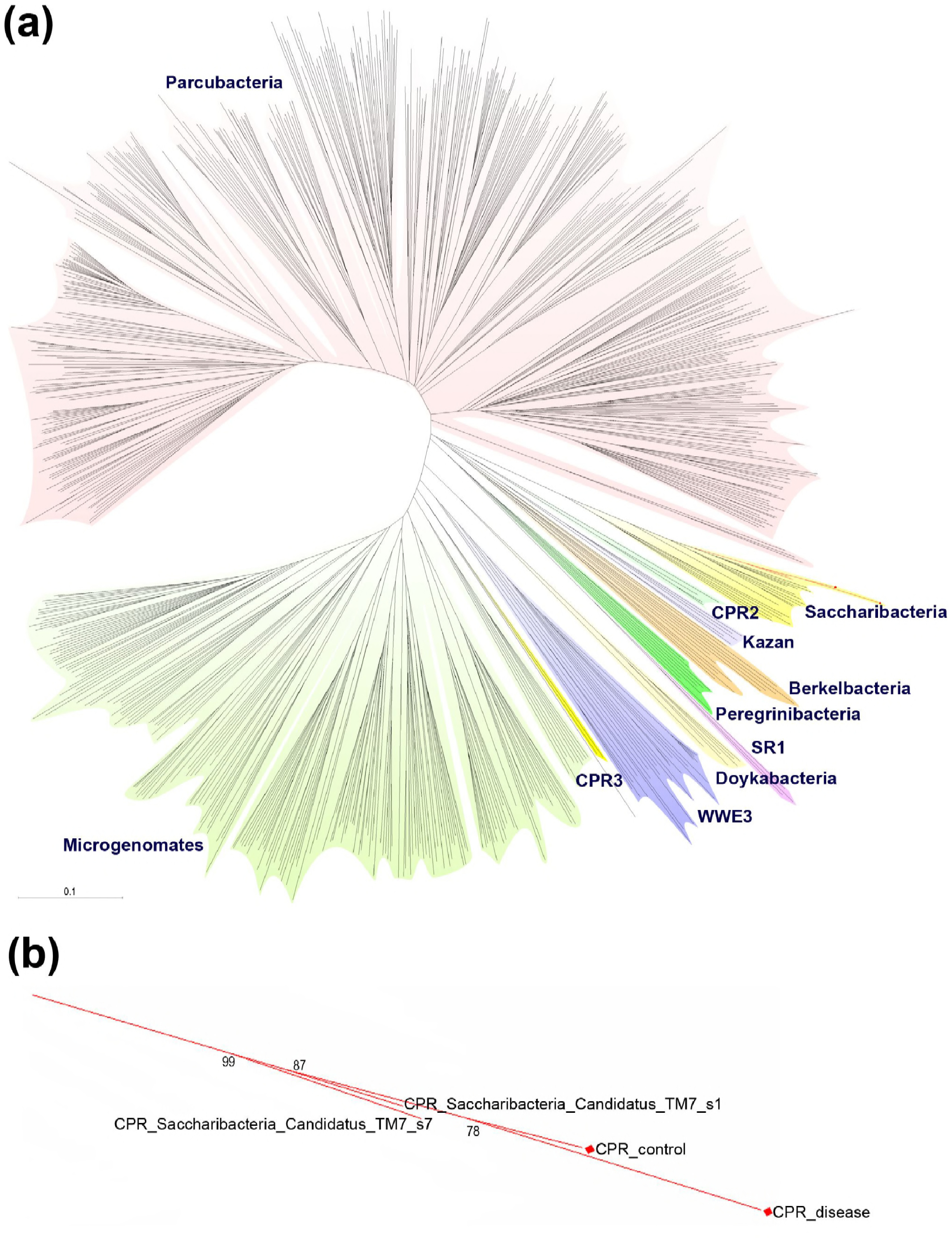
Phylogenetic tree of CPR generated from 16 concatenated ribosomal proteins. (a) The tree was constructed using Maximum Likelihood methods and visualized without detailed information for each organism. Each of the major lineages is assigned a color and is annotated with the phylum name. The clades of the newly identified CPR organisms and their closest relatives are indicated in red. (b) Magnified view of the clades containing the newly identified CPR organisms and their closest relatives. Bootstrap values (out of 100 replicates) are shown at the nodes.

In addition to the phylogenetic analyses described above, sequence similarity analyses (e.g., average nucleotide identity (ANI) comparisons) also suggested that the newly identified CPR genomes were closely related to the *Candidatus* TM7_s1 and *Candidatus* TM7_s7 organisms. ANI is the average nucleotide identity value calculated from a pair-wise comparison of homologous sequences between two genomes. Two genomes with an ANI value >95% can be regarded as members of the same species (26), and this metric is widely used to delineate species (26, 35). We calculated the ANI value between the newly constructed CPR genomes and some representative CPR genomes (**Table S2**). The largest ANI value for the CPR genome from the control group was 95.48% when compared with *Candidatus* TM7_s1. In addition, the largest ANI value for the CPR genome from the disease group was 95.76% to *Candidatus* TM7_s7. These results further suggest a close phylogenetic relationship of the genomes from the control and disease groups with *Candidatus* TM7_s1 and *Candidatus* TM7_s7, respectively.

We mapped all assembled contigs from our study to the CPR reference genome data set, which yielded best hits that were members of the TM7 phylum. The percentage of contigs that best matched genomes of the TM7 phylum were 70.2% and 79.7% in the control and disease groups, respectively, which represented much larger abundances than the organisms with the second most number of best matches. The reconstructed genome fragments of the control and disease groups aligned with high coverage to the reference genomes of *Candidatus* TM7_s1 and *Candidatus* TM7_s7 (**Fig. 3**). The correct assembly and classification of the newly constructed genomes were confirmed by direct visualization of the mapping of paired contigs.

**Fig 3.**
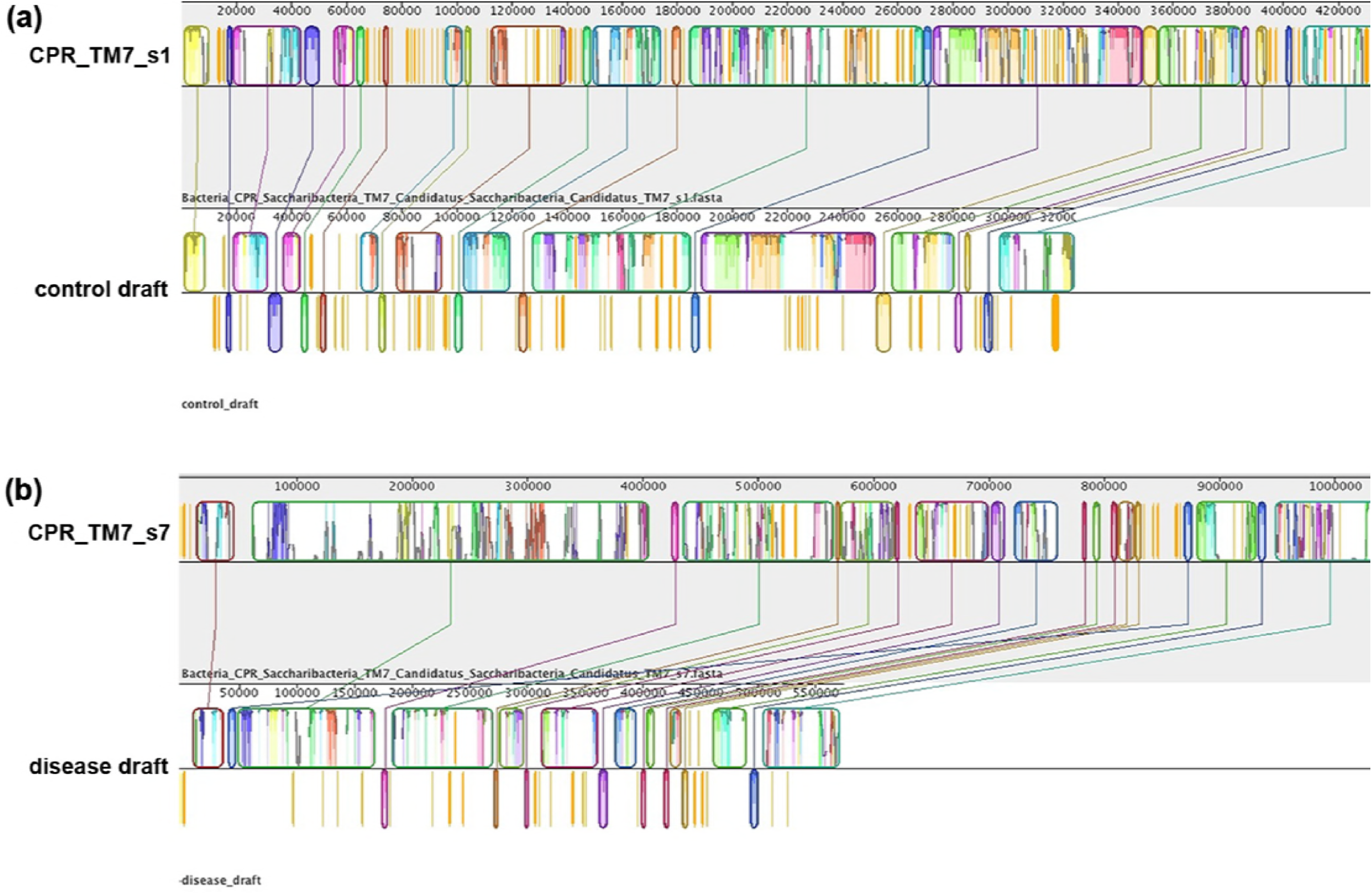
Comparisons of contig sequences of the two newly constructed CPR draft genomes and their closest relatives. (a) Sequence alignment of the CPR draft genome constructed in the control group metagenomes and its closest relative, *Candidatus* TM7_s1. (b) Sequence alignment of the CPR draft genome constructed from the disease group metagenomes and its closest relative, *Candidatus* TM7_s7. The alignments were produced using the MAUVE software package with default settings and using strains *Candidatus* TM7_s1 and *Candidatus* TM7_s7 as references.

The results described above indicate that both of the two newly constructed genomes derive from the TM7 superphylum. TM7 is a cosmopolitan and highly diverse division of bacteria, and were first identified in a German peat bog (36). TM7 can be found in diverse globally-distributed environments including soils, fresh waters, sea water, and hot springs (37). It has also been detected at a number of sites on human bodies including skin (38), the distal esophagus (39), the gut (6), and they are particularly prevalent in the oral cavity (40–45). It has been suggested that TM7 is a common, and perhaps permanent, component of the oral flora, and exhibits the capacity to maintain growth under both healthy and severe disease state conditions (40). The newly identified TM7 organisms in this study from healthy subjects and oral candidiasis patients were most closely related to *Candidatus* TM7_s1 and *Candidatus* TM7_s7, which together, are all found in human oral cavities.

TM7 have been predicted to exhibit potential pathogenic associations. Although TM7 were initially found in relatively low abundance in oral cavity samples (40), a more recent study demonstrated a high abundance of TM7 in subgingival plaque that was correlated with periodontal disease (46). Further, an increase in the proportion of TM7 in mild-periodontitis samples has been observed (40). In addition, 16S rRNA gene surveys have indicated a higher diversity of TM7 phylotypes in Crohn’s disease patients than in controls (6). Based on these findings, TM7 division species have been suggested to play an important role in the early stage of inflammatory mucosal processes, probably by modifying growth conditions for competing bacterial populations. Moreover, it has been hypothesized that TM7 could modulate the immune response of the host (11). However, as in our study, it is difficult to assess and compare physiological properties of these organisms in healthy and disease states, because they are difficult to cultivate and only a few culture representatives are known.

### The distribution of functional genes in the CPR genomes of healthy and disease patient groups

After identifying the affiliation of the CPR genomes in the control and disease groups with the TM7 superphylum, we investigated the functional predictions of proteins encoded in their genomes. Most of the functional genes within the two genomes encoded proteins related to cellular processes and signaling, in addition to hereditary information storage and processing. Cellular processes and signaling comprised 34% and 33% of the genes in the control and disease groups, respectively, suggesting an importance of signal transduction, cell motility and cell envelope biogenesis for these organisms. These observations are consistent with the phenotypic properties of TM7 isolates. A recent study demonstrated morphological changes in a TM7 cultivar from the human oral cavity that manifested as changes from ultrasmall cocci to elongated cells in response to environmental cues including oxygen levels and nutritional status (47). These cues could also involve activation of cellular signal transduction and cell motility pathways (**Fig. S3, Table S3**).

In contrast to other bacterial genomes (48), genes involved in metabolism comprised a small percentage of the newly constructed CPR genomes, with 27% and 24% in the control and disease groups, respectively. These results are congruent with the hypothesis of restricted metabolic capacities of CPR lineage organisms. Previous genomic analyses have indicated that the genomes of CPR organisms lack genes for the biosynthesis of most amino acids, nucleotides, vitamins, and lipids. Moreover, the genomes lack most of the genes for electron transport chain components and the tricarboxylic acid cycle. However, the metabolic abilities of some CPR organisms was recently expanded for members of the Parcubacteria, based on new genomes that encode putative components of the dissimilatory nitrate reduction to ammonia pathway (49, 50), as well as those involved in hydroxylamine oxidation (49). Metaproteomic analyses have also suggested that fermentative CPR likely play a significant role in hydrogen and carbon cycling in subsurface ecosystems (9, 51). In this study, we identified genes encoding various enzymes and transporters in the two newly constructed genomes, including the glycolysis-specific protein-coding genes for glyceraldehyde-3-phosphate dehydrogenase, triosephosphate isomerase, 6-phosphofructokinase, and enolase. Genes coding for the two former glycolysis proteins were found in both genomes, while the latter two were only identified in the genome from the disease group. Neither of the two reconstructed genomes had a complete tricarboxylic acid cycle pathway. These results are in accordance with previous hypotheses that CPR organisms may exhibit symbiotic lifestyles and are only capable of partial components of metabolic pathways. Nevertheless, the apparent lack of complete metabolic pathways could be due to incomplete genomes.

In conclusion, we designed a computational protocol to identify and characterize CPR organisms that are difficult to culture using metagenomic sequencing approaches. Specifically, phylogenetic analysis based on 16S rRNA genes, ribosomal proteins, and sequence similarity analyses (e.g., ANI comparisons and contig mapping) aid organism identification. Using this protocol, we reconstructed CPR genomes from human oral cavities with or without oral candidiasis using metagenomic data, and confirmed the affiliation of genomes. This culture-independent and amplicon-independent approach is a promising tool that can be widely used in the identification and characterization of various microorganisms, including CPR bacteria, fungi, viruses, and phages from oral cavity environments and elsewhere.

## Materials and Methods

### Sample collection

Samples were collected between March 2014 and April 2015 from outpatients aged 45–65 years that attended the department of oral mucosa at the Peking University School and Hospital of Stomatology, China. The study was approved by the ethics committee of the Peking University School and Hospital of Stomatology (Approval number PKUSSIRB-2013034). Saliva samples were collected only after informed written consent was obtained from all participants. The exclusion criteria were as follows: removable dentures; xerostomia (unstimulated salivary flow rate <1.0 ml/10 min); the use of steroid drugs, immunosuppressants, antibiotics, or mouthwash in the last 3 months; head or neck radiotherapy in the last 3 months; diabetes, cancer, anemia, or AIDS; severe local or systemic infections; and smoking history. Participants with clinical manifestations of oral candidiasis and a positive result from mycological examination (smear and culture) were included in the disease group (n = 12; median age: 57 years, range: 47–65 years; male/female: 5/7). Participants with no clinical manifestations of oral candidiasis or other oral mucosal diseases, in addition to a negative result from mycological examination (smear and culture) were included in the control group (n = 14; median age: 54 years, range: 45–65 years; male/female: 7/7).

The participants were asked to accumulate saliva in their mouths for at least 1 minute, followed by collection of saliva into a 50 ml sterile collection tube with a screw cap, and this was repeated several times to collect at least 2 ml of saliva. Saliva samples were placed on ice within 1 hour, subpackaged into 2 ml aliquots, and centrifuged at 12,000 rpm for 20 min at 4°C. Pelleted cell debris were stored at -80°C until DNA extraction.

### DNA extraction, sequencing, and processing

Genomic DNA extraction was performed using the FastDNA SPIN kit for Soil (MP Biomedicals, USA). Briefly, salivary cells were mixed with buffer and a lysing matrix (a mixture of ceramic and silica particles) and mechanically homogenized in the FastPrep Instrument (MP Biomedicals, USA) for 40 s at a speed setting of 6.0 in order to lyse microbial cells. DNA was then extracted following the manufacturer’s instructions with minimal modifications. Extracted DNA concentrations were determined using a Nanodrop 8000 spectrophotometer (Thermo Scientific, USA), and the quality of DNA extracts was evaluated by agarose gel electrophoresis.

Genomic DNA was sheared using the Biorupter ^®^ Pico sonication device (Diagenode, Belgium). DNA fragments of approximately 200 bp were selected by agarose gel electrophoresis. DNA libraries for shotgun sequencing on the Illumina platform were then constructed using the NEBNext Ultra DNA Library Prep Kit for Illumina (Illumina Inc, USA) according to the manufacturer’s instructions. Metagenomic sequencing of the 26 samples was performed on an Illumina HiSeq 2000 platform (Illumina Inc, USA) by generating 125 bp paired-end read libraries with insert sizes varying from 155–266 bp (mean 210 ± 35.5 bp).

Low-quality reads and reads that mapped to human DNA were removed from the raw data, and the remaining sequences were further quality-filtered. Briefly, any sequences with more than three ambiguous bases were removed. We screened reads for the minimum percentage of high-quality bases (Q30 >50%) and trimmed low-quality bases (<Q30) on the terminal end. Paired reads with at least one read that mapped to the human reference genome (GRCh37/hg19) were removed using the SOAP2 software (http://soap.genomics.org.cn) (settings: -m 100 -x 1000) (22).

### Metagenomic binning, annotation, and abundance calculations

To generate draft genomes, we first generated a data set comprising all publicly available CPR genomes from the Joint Genome Institute’s IMG-M database (img.jgi.doe.gov) (see **Table S4** for organism names and NCBI accession numbers of the genomes included). In order to identify CPR-specific reads, all trimmed reads from our libraries were mapped to the CPR genome data set using the BWA aligner (http://bio-bwa.sourceforge.net) (23). The coverage of each library and the coverage of each reference genome were calculated for each sample, and were compared between the disease and control groups.

Subsequently, the CPR reads from samples in the disease and control groups were merged separately, and assembled into contigs and scaffolds using the idba_ud assembler with default parameters (24). The assembled scaffolds were mapped to the CPR genome data set using BLASTn (-evalue 1e-5, nt identity >40%). Draft CPR genomes from the disease and control groups were then constructed based on the blast results. Open reading frames (ORFs) were predicted for the draft genomes using MetaProdigal (25) and then annotated by comparing predicted ORFs to the non-redundant (nr) protein sequence database and the Clusters of Orthologous Groups database (http://www.ncbi.nih.gov/COG/) using BLASTp (-evalue 1e-5, -max_hsps 1, aa identity >40%, and length coverage of gene >50%) to obtain best hit results. Lastly, pair-wise ANI values for pairs of newly assembled genomes and published CPR representative genomes were calculated using Jspecies (http://imedea.uib-csic.es/jspecies/) (26).

### Phylogenetic analysis of bins

16S rRNA genes were extracted from bins using the SSU-Align program with default parameters (27), and then aligned to two separate 16S rRNA gene databases (9, 28) using SINA on the SILVA server (29, 30). The 16S rRNA genes from disease and control group genomes, as well as from the representative CPR genomes were aligned in MUSCLE (31). Unaligned ends and regions with ≥95% of gaps were trimmed and the final alignment was used to generate a 16S rRNA phylogenetic tree using MEGA7 (https://www.megasoftware.net) (32).

In addition, a concatenated ribosomal protein alignment-based tree was generated from 16 ribosomal proteins (ribosomal proteins L2, L3, L4, L5, L6, L14, L15, L16, L18, L22, L24, S3, S8, S10, S17, S19). Briefly, each ribosomal protein sequence was extracted from the genome bins by identifying them with BLASTp against the nr database. Protein alignments were performed independently for each sequence dataset using MUSCLE with default parameters (31). The resulting alignments were subsequently trimmed to remove ambiguously aligned ends as well as columns comprising >95% gaps. Taxa were removed if their available sequence data represented less than 50% of the expected alignment columns. A maximum-likelihood tree from the ribosomal protein alignment was constructed using the MEGA7 software with 100 bootstraps. The computational protocol described above is shown in **Fig. 4**.

**Fig 4.**
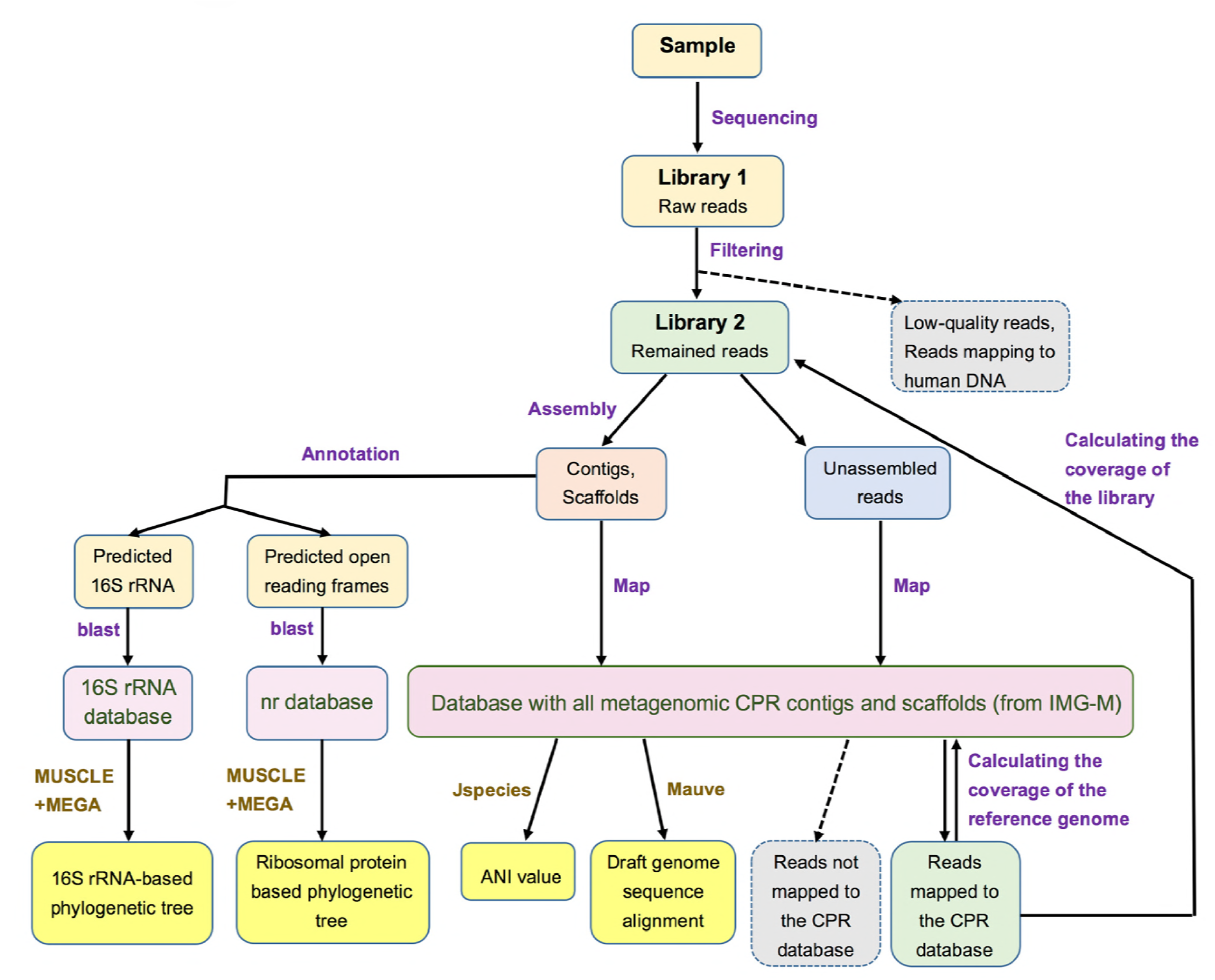
Overview of the computational workflow. Schematic of the protocol pipeline shows the steps required for detection of CPR organisms from metagenomic sequencing data. Boxes with pink backgrounds represent the databases used in the protocol, while yellow boxes represent four lines of evidence that demonstrate the affiliation of newly constructed genomes. Boxes with gray backgrounds and associated dotted arrows indicate sequence subsets that were discarded.

## Data availability

The metagenomic raw reads for 26 samples have been deposited in the sequence read archive under the project accession ID PRJNA478265.

## Acknowledgements

This work was financially supported by the National Natural Science Foundation of China (Nos. 81000441 and 81570985).

## Figure Legends

**Fig S1** Coverage of the sequencing library (a) and coverage of the reference genome (b) for reads mapped to CPR genomes in the reference dataset.

**Fig S2** An alternative visualization of the ribosomal protein tree in **Fig. 2** in which major lineages are annotated with an identical background as in **Fig. 2**.

**Fig S3** Venn diagram indicating the number of CPR genomes in the reference dataset that mapped to the contigs of the two newly constructed CPR draft genomes. Pie charts indicate COG functional classifications of the genes that were predicted in the two newly constructed genomes.

**Table S1** Information on the metagenomic sequencing libraries of 26 oral cavity samples.

**Table S2** ANI values calculated by cross-genome comparisons of the two newly constructed CPR draft genomes and the CPR species most closely related to them.

**Table S3** Detailed information of the COG functional classifications and annotations of genes predicted in the newly constructed CPR draft genomes from the control group (a) and the disease group (b).

**Table S4** NCBI or JGI IMG accession numbers for all genomes used in the self-constructed CPR dataset.

## References

1. Kolenbrander PE, Palmer RJ Jr, Periasamy S, Jakubovics NS. 2010. Oral multispecies biofilm development and the key role of cell-cell distance. Nat Rev Microbiol 8:471–480.

2. Zhang X, Zhang D, Jia H, Feng Q, Wang D, Liang D, Wu X, Li J, Tang L, Li Y, Lan Z, Chen B, Li Y, Zhong H, Xie H, Jie Z, Chen W, Tang S, Xu X, Wang X, Cai X, Liu S, Xia Y, Li J, Qiao X, Al-Aama JY, Chen H, Wang L, Wu QJ, Zhang F, Zheng W, Li Y, Zhang M, Luo G, Xue W, Xiao L, Li J, Chen W, Xu X, Yin Y, Yang H, Wang J, Kristiansen K, Liu L, Li T, Huang Q, Li Y, Wang J. 2015. The oral and gut microbiomes are perturbed in rheumatoid arthritis and partly normalized after treatment. Nat Med 21:895–905.

3. Sardi JC, Duque C, Camargo GA, Hofling JF, Gonçalves RB. 2011. Periodontal conditions and prevalence of putative periodontopathogens and Candida spp. in insulin-dependent type 2 diabetic and non-diabetic patients with chronic periodontitis–a pilot study. Arch Oral Biol 56:1098–1105.

4. Kong EF, Kucharíková S, Van Dijck P, Peters BM, Shirtliff ME, Jabra-Rizk MA. 2015. Clinical implications of oral candidiasis: host tissue damage and disseminated bacterial disease. Infect Immun 83:604–613.

5. Kianoush N, Adler CJ, Nguyen KA, Browne GV, Simonian M, Hunter N. 2014. Bacterial profile of dentine caries and the impact of pH on bacterial population diversity. PLoS One 9:e92940.

6. Kuehbacher T, Rehman A, Lepage P, Hellmig S, Fölsch UR, Schreiber S, Ott SJ. 2008. Intestinal TM7 bacterial phylogenies in active inflammatory bowel disease. J Med Microbiol 57:1569–1576.

7. Paster BJ, Boches SK, Galvin JL, Ericson RE, Lau CN, Levanos VA, Sahasrabudhe A, Dewhirst FE. 2001. Bacterial diversity in human subgingival plaque. J Bacteriol 183:3770–3783.

8. Soro V, Dutton LC, Sprague SV, Nobbs AH, Ireland AJ, Sandy JR, Jepson MA, Micaroni M, Splatt PR, Dymock D, Jenkinson HF. 2014. Axenic culture of a candidate division TM7 bacterium from the human oral cavity and biofilm interactions with other oral bacteria. Appl Environ Microbiol 80:6480–6489.

9. Brown CT, Hug LA, Thomas BC, Sharon I, Castelle CJ, Singh A, Wilkins MJ, Wrighton KC, Williams KH, Banfield JF. 2015. Unusual biology across a group comprising more than 15% of domain Bacteria. Nature 523:208–211.

10. Campbell JH, O’Donoghue P, Campbell AG, Schwientek P, Sczyrba A, Woyke T, Söll D, Podar M. 2013. UGA is an additional glycine codon in uncultured SR1 bacteria from the human microbiota. Proc Natl Acad Sci U S A 110:5540–5545.

11. He X, McLean JS, Edlund A, Yooseph S, Hall AP, Liu SY, Dorrestein PC, Esquenazi E, Hunter RC, Cheng G, Nelson KE, Lux R, Shi W. 2015. Cultivation of a human-associated TM7 phylotype reveals a reduced genome and epibiotic parasitic lifestyle. Proc Natl Acad Sci U S A 112:244–249.

12. Luef B, Frischkorn KR, Wrighton KC, Holman HY, Birarda G, Thomas BC, Singh A, Williams KH, Siegerist CE, Tringe SG, Downing KH, Comolli LR, Banfield JF. 2015. Diverse uncultivated ultra-small bacterial cells in groundwater. Nat Commun 6:6372.

13. Rinke C, Schwientek P, Sczyrba A, Ivanova NN, Anderson IJ, Cheng JF, Darling A, Malfatti S, Swan BK, Gies EA, Dodsworth JA, Hedlund BP, Tsiamis G, Sievert SM, Liu WT, Eisen JA, Hallam SJ, Kyrpides NC, Stepanauskas R, Rubin EM, Hugenholtz P, Woyke T. 2013. Insights into the phylogeny and coding potential of microbial dark matter. Nature 499:431–437.

14. Solden L, Lloyd K, Wrighton K. 2016. The bright side of microbial dark matter: lessons learned from the uncultivated majority. Curr Opin Microbiol 31:217–226.

15. Burstein D, Sun CL, Brown CT, Sharon I, Anantharaman K, Probst AJ, Thomas BC, Banfield JF. 2016. Major bacterial lineages are essentially devoid of CRISPR-Cas viral defence systems. Nat Commun 7:10613.

16. Gong J, Qing Y, Guo X, Warren A. 2014. “Candidatus Sonnebornia yantaiensis”, a member of candidate division OD1, as intracellular bacteria of the ciliated protist Paramecium bursaria (Ciliophora, Oligohymenophorea). Syst Appl Microbiol 37:35–41.

17. Kantor RS, Wrighton KC, Handley KM, Sharon I, Hug LA, Castelle CJ, Thomas BC, Banfield JF. 2013. Small genomes and sparse metabolisms of sediment-associated bacteria from four candidate phyla. MBio 4:e00708–13.

18. Nelson WC, Stegen JC. 2015. The reduced genomes of Parcubacteria (OD1) contain signatures of a symbiotic lifestyle. Front Microbiol 6:713.

19. Danczak RE, Johnston MD, Kenah C, Slattery M, Wrighton KC, Wilkins MJ. 2017. Members of the Candidate Phyla Radiation are functionally differentiated by carbon- and nitrogen-cycling capabilities. Microbiome 5:112.

20. Klindworth A, Pruesse E, Schweer T, Peplies J, Quast C, Horn M, Glöckner FO. 2013. Evaluation of general 16S ribosomal RNA gene PCR primers for classical and next-generation sequencing-based diversity studies. Nucleic Acids Res 41:e1.

21. Dick GJ, Andersson AF, Baker BJ, Simmons SL, Thomas BC, Yelton AP, Banfield JF. 2009. Community-wide analysis of microbial genome sequence signatures. Genome Biol 10:R85.

22. Li R, Yu C, Li Y, Lam TW, Yiu SM, Kristiansen K, Wang J. 2009. SOAP2: an improved ultrafast tool for short read alignment. Bioinformatics 25:1966–1967.

23. Li H, Durbin R. 2009. Fast and accurate short read alignment with Burrows-Wheeler transform. Bioinformatics 25:1754–1760.

24. Peng Y, Leung HC, Yiu SM, Chin FY. 2012. IDBA-UD: a de novo assembler for single-cell and metagenomic sequencing data with highly uneven depth. Bioinformatics 28:1420–1428.

25. Hyatt D, Chen GL, Locascio PF, Land ML, Larimer FW, Hauser LJ. 2010. Prodigal: prokaryotic gene recognition and translation initiation site identification. BMC Bioinformatics 11:119.

26. Goris J, Konstantinidis KT, Klappenbach JA, Coenye T, Vandamme P, Tiedje JM. 2007. DNA-DNA hybridization values and their relationship to whole-genome sequence similarities. Int J Syst Evol Microbiol 57:81–91.

27. Nawrocki EP. 2009. Structural RNA homology search and alignment using covariance models. Dissertation, University of Washington.

28. Anantharaman K, Brown CT, Hug LA, Sharon I, Castelle CJ, Probst AJ, Thomas BC, Singh A, Wilkins MJ, Karaoz U, Brodie EL, Williams KH, Hubbard SS, Banfield JF. 2016. Thousands of microbial genomes shed light on interconnected biogeochemical processes in an aquifer system. Nat Commun 7:13219.

29. Pruesse E, Peplies J, Glöckner FO. 2012. SINA: accurate high-throughput multiple sequence alignment of ribosomal RNA genes. Bioinformatics 28:1823–1829.

30. Quast C, Pruesse E, Yilmaz P, Gerken J, Schweer T, Yarza P, Peplies J, Glöckner FO. 2013. The SILVA ribosomal RNA gene database project: improved data processing and web-based tools. Nucleic Acids Res 41:D590–596.

31. Edgar RC. 2004. MUSCLE: multiple sequence alignment with high accuracy and high throughput. Nucleic Acids Res 32:1792–1797.

32. Kumar S, Stecher G, Tamura K. 2016. MEGA7: Molecular Evolutionary Genetics Analysis version 7.0 for bigger datasets. Mol Biol Evol 33:1870.

33. Hug LA, Baker BJ, Anantharaman K, Brown CT, Probst AJ, Castelle CJ, Butterfield CN, Hernsdorf AW, Amano Y, Ise K, Suzuki Y, Dudek N, Relman DA, Finstad KM, Amundson R, Thomas BC, Banfield JF. 2016. A new view of the tree of life. Nat Microbiol 1:16048.

34. Hug LA, Castelle CJ, Wrighton KC, Thomas BC, Sharon I, Frischkorn KR, Williams KH, Tringe SG, Banfield JF. 2013. Community genomic analyses constrain the distribution of metabolic traits across the Chloroflexi phylum and indicate roles in sediment carbon cycling. Microbiome 1:22.

35. Chan JZ, Halachev MR, Loman NJ, Constantinidou C, Pallen MJ. 2012. Defining bacterial species in the genomic era: insights from the genus Acinetobacter. BMC Microbiol 12:302.

36. Rheims H, Spröer C, Rainey FA, Stackebrandt E. 1996. Molecular biological evidence for the occurrence of uncultured members of the actinomycete line of descent in different environments and geographical locations. Microbiology 142:2863–2870.

37. Hugenholtz P, Tyson GW, Webb RI, Wagner AM, Blackall LL. 2001. Investigation of candidate division TM7, a recently recognized major lineage of the domain Bacteria with no known pure-culture representatives. Appl Environ Microbiol 67:411–419.

38. Dinis JM, Barton DE, Ghadiri J, Surendar D, Reddy K, Velasquez F, Chaffee CL, Lee MC, Gavrilova H, Ozuna H, Smits SA, Ouverney CC. 2011. In search of an uncultured human-associated TM7 bacterium in the environment. PLoS One 6:e21280.

39. Pei Z, Bini EJ, Yang L, Zhou M, Francois F, Blaser MJ. 2004. Bacterial biota in the human distal esophagus. Proc Natl Acad Sci U S A 101:4250–4255.

40. Brinig MM, Lepp PW, Ouverney CC, Armitage GC, Relman DA. 2003. Prevalence of bacteria of division TM7 in human subgingival plaque and their association with disease. Appl Environ Microbiol 69:1687–1694.

41. Colombo AP, Boches SK, Cotton SL, Goodson JM, Kent R, Haffajee AD, Socransky SS, Hasturk H, Van Dyke TE, Dewhirst F, Paster BJ. 2009. Comparisons of subgingival microbial profiles of refractory periodontitis, severe periodontitis, and periodontal health using the human oral microbe identification microarray. J Periodontol 80:1421–1432.

42. Duran-Pinedo AE, Chen T, Teles R, Starr JR, Wang X, Krishnan K, Frias-Lopez J. 2014. Community-wide transcriptome of the oral microbiome in subjects with and without periodontitis. ISME J 8:1659–1672.

43. Marcy Y, Ouverney C, Bik EM, Lösekann T, Ivanova N, Martin HG, Szeto E, Platt D, Hugenholtz P, Relman DA, Quake SR. 2007. Dissecting biological “dark matter” with single-cell genetic analysis of rare and uncultivated TM7 microbes from the human mouth. Proc Natl Acad Sci U S A 104:11889–11894.

44. Rego RO, Oliverira CA, dos Santos-Pinto A, Jordan SF, Zambon JJ, Cirelli JA, Haraszthy VI. 2010. Clinical and microbiological studies of children and adolescents receiving orthodontic treatment. Am J Dent 23:317–323.

45. Teles FR, Teles RP, Siegelin Y, Paster B, Haffajee AD, Socransky SS. 2011. RNA-oligonucleotide quantification technique (ROQT) for the enumeration of uncultivated bacterial species in subgingival biofilms. Mol Oral Microbiol 26:127–139.

46. Liu B, Faller LL, Kiltgord N, Mazumdar V, Ghodsi M, Sommer DD, Gibbons TR, Treangen TJ, Chang YC, Li S, Stine OC, Hasturk H, Kasif S, Segrè D, Pop M, Amar S. 2012. Deep sequencing of the oral microbiome reveals signatures of periodontal disease. PLoS One 7:e37919.

47. Bor B, Poweleit N, Bois JS, Cen L, Bedree JK, Zhou ZH, Gunsalus RP, Lux R, McLean JS, He X, Shi W. 2016. Phenotypic and physiological characterization of the epibiotic interaction between TM7x and its basibiont Actinomyces. Microb Ecol 71:243–255.

48. Meng P, Lu C, Zhang Q, Lin J, Chen F. 2017. Exploring the genomic diversity and cariogenic differences of Streptococcus mutans strains through pan-genome and comparative genome analysis. Curr Microbiol 74:1200–1209.

49. Castelle CJ, Brown CT, Thomas BC, Williams KH, Banfield JF. 2017. Unusual respiratory capacity and nitrogen metabolism in a Parcubacterium (OD1) of the Candidate Phyla Radiation. Sci Rep 7:40101.

50. León-Zayas R, Peoples L, Biddle JF, Podell S, Novotny M, Cameron J, Lasken RS, Bartlett DH. 2017. The metabolic potential of the single cell genomes obtained from the Challenger Deep, Mariana Trench within the candidate superphylum Parcubacteria (OD1). Environ Microbiol 19:2769–2784.

51. Wrighton KC, Castelle CJ, Wilkins MJ, Hug LA, Sharon I, Thomas BC, Handley KM, Mullin SW, Nicora CD, Singh A, Lipton MS, Long PE, Williams KH, Banfield JF. 2014. Metabolic interdependencies between phylogenetically novel fermenters and respiratory organisms in an unconfined aquifer. ISME J 8:1452–1463.

